# IFNB induced by non-lytic virus immunotherapy promotes improved survival in hepatocellular carcinoma, mediated by MHCII-independent cytotoxic CD4^+^ T-cells

**DOI:** 10.1101/2024.02.23.581738

**Authors:** Russell Hughes, Amy Moran, Samantha Garcia-Cardenas, Karen J. Scott, Elizabeth Appleton, Matthew J. Bentham, Elizabeth Ilett, Alan Melcher, Andrew Macdonald, Adel Samson, Stephen Griffin

## Abstract

Hepatocellular carcinoma (HCC) is the third most common cause of cancer deaths worldwide. Combination immunotherapy is now standard of care for advanced HCC, improving patient outcomes. However, a considerable number of patients remain unresponsive, or are unable to tolerate therapy. Tyrosine kinase inhibitors (TKIs), such as the former first-line agent sorafenib, remain an option for such patients, yet provide only marginal efficacy. We hypothesised that a clinically advanced immunogenic “oncolytic” virus, namely, human *Orthoreovirus*, might improve TKI mediated therapy. Surprisingly, *uv*-inactivated, replication-deficient reovirus, but not live virus, significantly extended survival when combined with sorafenib in preclinical immunocompetent HCC models. Favourable responses were dependent upon adaptive immunity, mediated by IFNB-induced skewing of the infiltrating T-cell ratio in favour of cytotoxic CD4^+^ T-cells expressing granzyme B and perforin. Interestingly, this subset effectively killed tumours via both contact juxtacrine and paracrine processes, the former being MHCII independent. Moreover, efficacy correlated with more rapid and robust IFN production by inactivated virus due to the absence of innate viral antagonists. Thus, we reveal a means to improve TKI-HCC outcomes through an alternative virus-driven immunotherapy, underpinned by non-classical immunological mechanisms.

**IMPACT AND IMPLICATIONS:** Immune checkpoint immunotherapy is revolutionising cancer treatment, yet considerable numbers of patients still fail to respond and must resort to older, more toxic and less effective therapies, including Sorafenib for the management of HCC.

We demonstrate that burgeoning virus-driven immunotherapy can be successfully combined with Sorafenib to extend preclinical HCC survival, but only when the virus is *uv*-inactivated to prevent already attenuated innate immune antagonism, specifically increasing the magnitude of tumour IFNB responses. IFNB was essential to promote tumour infiltration of cytotoxic CD4+ cells during therapy, which was a hallmark of long-term survival mediated by ensuing adaptive responses.

We anticipate this work will be of interest to clinicians and cancer immunology researchers, promoting closer inspection of the immune microenvironment and cancer-specific responses to OV therapy, specifically those driven by non-canonical anti-cancer mechanisms involving IFNB and cytotoxic CD4*+* T-cells.

## INTRODUCTION

Hepatocellular carcinoma (HCC) is the third leading cause of cancer deaths worldwide. A significant proportion of HCC patients (∼40%) present with advanced disease at diagnosis, excluding them from potentially curative surgery. Consequently, ∼88% of patients succumb within 5-years post-diagnosis across disease stages. Recently, immunotherapy combinations, including licensed treatments targeting PD-L1 and VEGF-A have dramatically improved clinical outcomes, dependent upon a favourable immunological tumour microenvironment. Patients failing to respond revert to prior systemic TKI therapy, most commonly sorafenib. The survival benefit gained following sorafenib treatment is limited^1^, as is patient compliance. Thus, considerable need exists for novel therapeutic approaches that augment existing TKI efficacy.

Oncolytic viruses (OV) are a promising area of cancer immunotherapy and are well tolerated by patients. Numerous OVs are in clinical trials for a range of solid and hematological malignancies. OV-therapies are known to exert their anti-cancer activity by both direct tumour cell lysis and stimulation of anti-tumour immunity. This arises due to innate responses, direct leukocyte stimulation, and the liberation of PAMPs, DAMPs and other cell components including tumour antigens. The balance between how these processes combine to achieve efficacy logically varies according to both the OV, as well as the tumour in question.

Oncolytic human *Orthoreovirus* (type-3, Dearing Strain, supplied as Pelareorep, Oncolytics Biotech, Calgary, CA. Referred to herein as “Reo”) has been both widely and safely used in numerous cancer clinical trials and is currently under fast-track review for both metastatic breast cancer and advanced/metastatic pancreatic cancer^2^. Although Reo exerts direct lytic effects following replication in cancerous cells, there is considerable interest in its pronounced ability to activate the immune system^3–7^. The segmented, dsRNA genome of Reovirus, including its terminal diphosphate, can be detected by innate cellular pattern recognition receptors (PRR) including endosomal Toll-like receptors (TLR3) and cytosolic RIG-like receptors^8^ (RIG-I and MDA-5). Upon activation, PRRs trigger signalling cascades that drive host cells to produce a wide variety of inflammatory cytokines, particularly interferons. These play a major role inducing an anti-viral state by upregulating the expression of interferon-stimulated genes (ISGs), but also stimulate and enhance immune cell function. In the context of cancer, Reo stimulates antigen-specific cytotoxic CD8^+^ T-cell responses^4–7^ capable of breaking tumour immune tolerance in pre-clinical mouse models^5, 6^. Immune activation is also observed in Reo treated patients, evidenced by increased expression of the interferon gamma-inducible immune checkpoint molecule, PD-L1^9^, increased levels of anti-Reo neutralising antibodies in peripheral blood^10, 11^, and the accumulation of intra-tumoural T-cells^12^.

CD4^+^ T-helper cells (T_H_-cells) are critical master co-ordinators of the adaptive immune system. In addition, under specific conditions T_H_-cells exert direct toxicity against a range of cell types. Cytotoxic CD4^+^ T_H_-cells (CTHs) are frequently observed in patients with chronic viral infections, including both human cytomegalovirus (hCMV)^13^ and human immunodeficiency virus (HIV)^14^, as well as in pre-clinical models of lymphocytic choriomeningitis (LCMV) ^15^.

CTHs resemble *bona fide* cytotoxic CD8^+^ T-cells (CTLs) by expressing granzyme-B and perforin, although rather than MHCI, they exert MHCII-dependent, antigen-specific cell killing. T_H_-cells also deploy granzyme/perforin-independent cytotoxicity, including FasL/Fas and TNF-related apoptosis-inducing ligand (TRAIL), which are involved in maintenance of peripheral tolerance through activation-induced cell death (AICD), and the elimination of malignant or virus-infected cells^16, 17^.

Here, we describe a striking observation whereby *uv*-inactivated, replication-deficient Reo (*uv*-Reo), but *not* live Reo, dramatically extended survival in a pre-clinical, syngeneic, immunocompetent HCC model when combined with sorafenib. Whilst long-term protection relied upon adaptive responses, its inception was critically dependent upon expression of IFNB, and a T_H_1-dominated anti-tumour immune response underpinned, surprisingly, by multiple MHCII-*independent* modes of killing. Mechanistically, tumour-borne expression of the T_H_1-cell tropic chemokine, CCL5, led to a markedly increased CD4:CD8 ratio amongst tumour-infiltrating lymphocytes (TIL) during therapy. *uv*-Reo/sorafenib induced CTH exhibited both paracrine as well as proximity-dependent tumour cell killing. Sorafenib sensitised tumour cells to secreted TNFA and IFNG produced by CTHs whereas IFNB engaged a proximity-dependent mode of killing that was reliant on granzyme B and perforin, but *not* MHCII. The superiority of the *uv*-Reo response was attributable to the absence of viral IFN antagonists, raising questions over the possible limitations of other, less attenuated, virus-driven immunotherapies.

## MATERIALS & METHODS

### Mouse models

All *in vivo* experiments were conducted with the approval of the University of Leeds Applications and Amendments (Ethics) Committee and in accordance with UK Home Office regulations (PP1816772). BALB/c, SCID, and SCID/Beige mice were housed in isolator cages with 12-h light/dark cycles at 22°C with access to food and water *ad libitum*. For overall survival studies, female mice, aged 7 – 8 weeks, were implanted subcutaneously (*s.c.,*) with 1MEA cells in 100 *µ*L of PBS. Once palpable (∼2 – 3 mm in diameter), mice were treated with either Reo or *uv*-Reo (1x10^7^ pfu) *via* intra-tumoural injection (*i.t.,*), three times per week for six weeks. Sorafenib (10 mg/Kg) or vehicle (PBS, 25% PEG-400, 5% Tween-20, 5% ethanol) were administered by oral gavage (*o.g.,*), daily for 4 four weeks. Tumour diameter was measured in two dimensions daily and mice were culled when they reached 15 mm in diameter as a proxy for cancer-induced mortality. For histological, proteomic and RNA analyses, female mice were implanted with murine HCC cells and treated with live- or *uv*-Reo alone or in combination with sorafenib or vehicle as described above, for two weeks. Twenty-four hours after the last treatment the mice were culled by an approved Schedule 1 method and tissue processed accordingly.

### Primary cell cultures and cell lines

Human (HLE) and murine (1ME.A7.7R.1 [1MEA]) HCC cell lines, and primary immune cells, were maintained in humidified incubators with 5% CO_2_ in DMEM supplemented with 2 mM *l*-glutamine, 10% FBS, and 1% non-essential amino acids. Human and murine T-cells were isolated by a combination of positive selection using antibody-conjugated magnetic beads directed against CD4 and CD8 for whole blood (Human) or negative selection from spleens and lymph nodes (Mouse). Activation of human and murine T-cells was performed using plate-bound anti-CD3 and medium supplemented with anti-CD28 for three days, with or without recombinant IL2 and IL12 for T_H_1 polarisation.

### Chemotaxis assay

CD4^+^ T-cells were serum-starved for two hours in chemotaxis buffer (RPMI-1640 and 0.5 % BSA) prior to assay. For each experiment, 3x10^5^ CD4^+^ T-cells, labelled with CFSE, were seeded into tissue culture inserts (5 *µ*m pore size), in 24-well plates, in chemotaxis buffer in the presence or absence a CCR5 inhibitor (Maraviroc, 1 *µ*M) or vehicle. The lower compartment contained either chemotaxis buffer alone or was supplemented with CCL5 (100 ng/mL) with or without Maraviroc or vehicle. Chemotaxis assays were stopped after 1.5 hours and cells counted using flow cytometry.

### Immunofluorescence

Tumour cryosections 14 *µ*m thick were fixed with either ice-cold acetone or 4% paraformaldehyde, blocked in appropriate serum and incubated with fluorophore-conjugated primary antibodies (2 – 5 *µ*g/mL) in PBS, overnight at 4°C. Nuclei were counter-stained using DAPI and labelled sections were mounted in ProLong Diamond antifade reagent. Random fields of view (F.O.V.) were acquired using both a Nikon A1R and Zeiss LSM 980 confocal microscopes. Image processing and quantification was performed using ‘Fiji’ Image.

### Proteomics

Cytokine arrays were performed using tumour protein extracts generated from snap-frozen biopsies using a combination of bead mill and freeze/fracture in PBS. Mouse cytokine arrays were incubated with 1 mg pooled total protein. Membranes were developed using Pierce^TM^ chemiluminescent substrate with a ChemiDoc^TM^ imaging system and image quantification was performed using ‘Fiji’ Image J. For ELISAs, clarified supernatants were generated from human and mouse HCC cell lines following incubation overnight with Reo/*uv*-Reo (2 PFU/cell) in the presence or absence of sorafenib (7µM), and from T-cells at three days post-activation.

### Flow cytometry

Antibody labelling of T-cells and HCC cell lines was performed in staining buffer (HBSS + 0.5% BSA), on ice, using directly-conjugated antibodies (1 – 5 µg/mL). For cryopreserved tumour biopsies, single cell suspensions were generated by passing tumours through a 70 µm nylon mesh with subsequent labelling of both cell surface and intra-cellular antigens using a CytoFix/CytoPerm Kit. Data were collected using a CytoFLEX S flow cytometer.

### HCC/T-cell co-culture killing assays

For direct co-culture killing assays, human and murine CFSE-labelled HCC target cells (1.5x10^4^) were co-cultured with near infra-red dye-labelled T-cells at a ratio of 50:1 in the presence or absence of IFNB, with or without neutralising antibodies, EGTA, GZMB inhibitor (z-AAD-CMK) or Caspase inhibitor (z-VAD-FMK), where indicated. Following overnight incubation, all cells were collected and stained with Zombie UV viability dye then fixed in 4% PFA prior to analysis. Target cell killing (CFSE^+^NIR^neg^) was determined using a CytoFLEX S flow cytometer. For indirect co-culture killing assays, T-cells and target cells were separated using tissue culture inserts with a 0.4 µm pore size.

### RNASeq and immune deconvolution

RNA samples were extracted from 1MEA tumours using an RNeasy mini kit and subjected to Illumina sequencing (Novogene UK Ltd), with a sequencing depth of 20 million reads. RNASeq data were uploaded to the TIMER2.0 online immune estimation resource and the xCell immune deconvolution algorithm was applied^18–20^.

### Statistics

All figures and statistical analyses were performed using Prism software (GraphPad, San Diego, CA). All data presented are expressed as means ± standard error of the mean (SEM) and were analysed by one-tailed or two-tailed unpaired Student’s t-test where appropriate. P values less than 0.05 were considered statistically significant and marked as follows; * p<0.05, ≠ p<0.01, + <0.001, ^ <0.0001. Sample size (n) is indicated where appropriate in figure legends.

## RESULTS

### Suppression of HCC tumour growth during *uv*-Reo/sorafenib therapy involves the skewing of intra-tumoural T-cell ratios (T_H_:CTL) in favour of CD4^+^ T_H_-cells

We investigated the therapeutic effect of combining sorafenib with Reo in Balb/c mice bearing syngeneic, subcutaneous 1MEA HCC tumours, controlling for virus gene expression-mediated effects by including *uv*-inactivated, replication-deficient virus (*uv*-Reo - **Fig. S1A**-**B**). Surprisingly, the combination of *uv*-Reo and sorafenib significantly extended the survival of tumour-bearing mice well beyond that of all other treatment groups (**Fig. 1A**), a response that was greatly reduced in SCID mice and lost entirely in SCID/Beige, supporting an immune-mediated mechanism significantly driven by the adaptive response (**Fig. 1B**).

**Figure 1.**
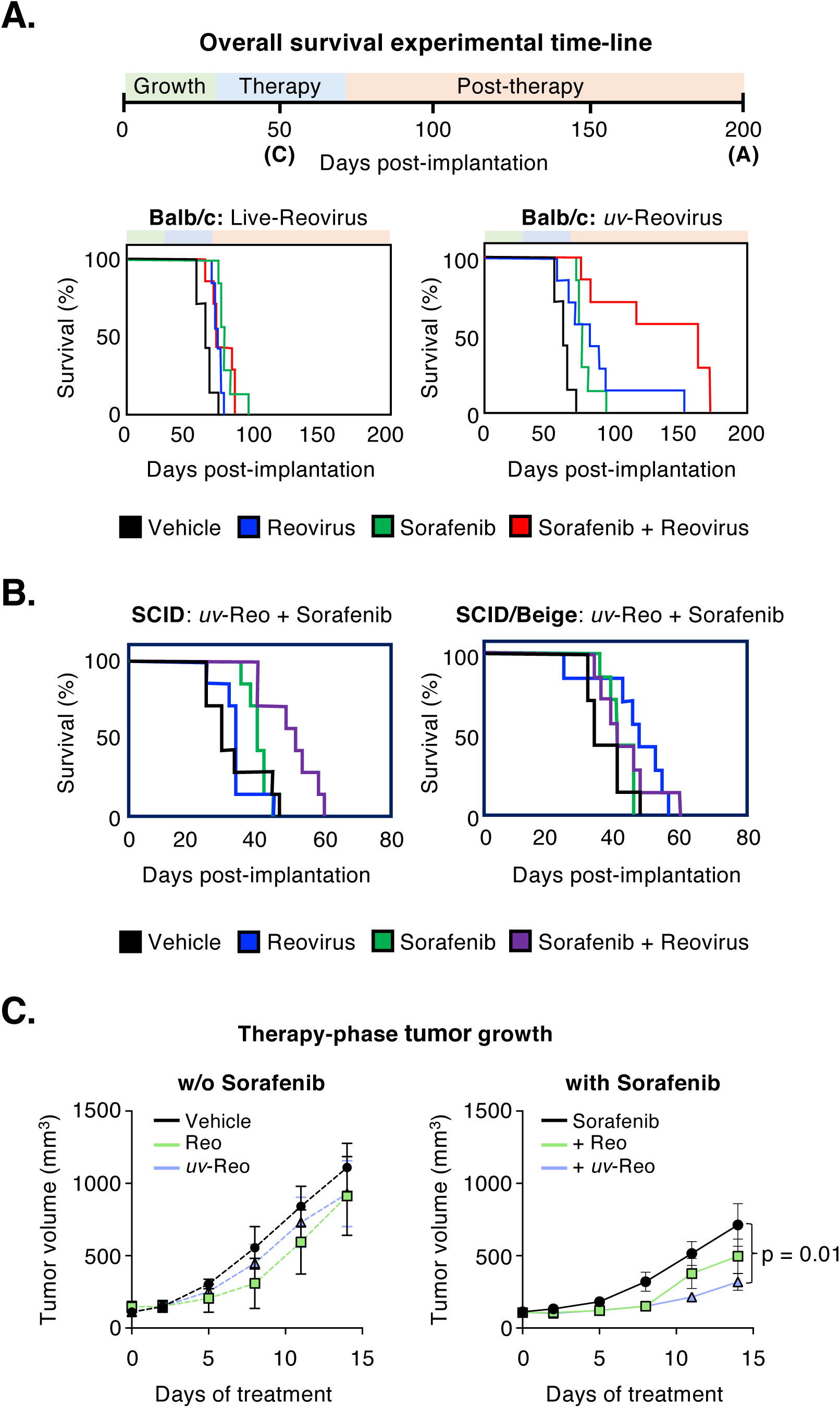
The survival of mice bearing syngeneic hepatocellular carcinomas is significantly extended by an immune-mediated mechanism induced by a combination of sorafenib and *uv*-Reo. (**A**) Overall survival analysis of 1MEA tumour-bearing mice treated with Reo (*left*) or uv-Reo (*right*) alone or in combination with sorafenib. All treatments were significant relative to vehicle (p≤0.05) and *uv*-Reo/Sorafenib was significant compared to *uv*-Reo (p<0.0189) and sorafenib (p<0.0027) monotherapies. (**B**) Overall survival analysis of SCID (*left*) and SCID/Beige (*right*) mice bearing 1MEA tumours treated as with *uv*-Reo alone or in combination with sorafenib. *uv*-Reo/sorafenib was significant compared to all groups in SCID mice only (p≤0.01). (**C**) Tumour volumetric data from immunocompetent Balb/c mice bearing 1MEA tumours treated with Reo/*uv*-Reo monotherapy or in combination with sorafenib and culled after two-weeks of therapy for IF analysis (n = 5 mice per condition).

Thus, we examined the leukocyte composition of 1MEA tumours, grown in Balb/c hosts, harvested when responding to therapy (Therapy phase – **Fig 1A** & **Fig. 1C**), to identify changes in infiltrating immune cell(s) that contribute to the subsequent improved survival phenotype. The only leukocyte population found to be increased in tumours treated with *uv*-Reo/sorafenib combination therapy relative to control animals was the CD3^+^CD4^+^ T_H_-cell, but not CD3^+^CD8^+^ CTLs (**Fig. 2A – C** & **Fig. S2**). This significantly skewed the T-cell ratio in favour of CD4^+^ cells (**Fig. 2D**) implying that the response to therapy that underpinned the ensuing improved survival was dominated by T_H_-cells.

**Figure 2.**
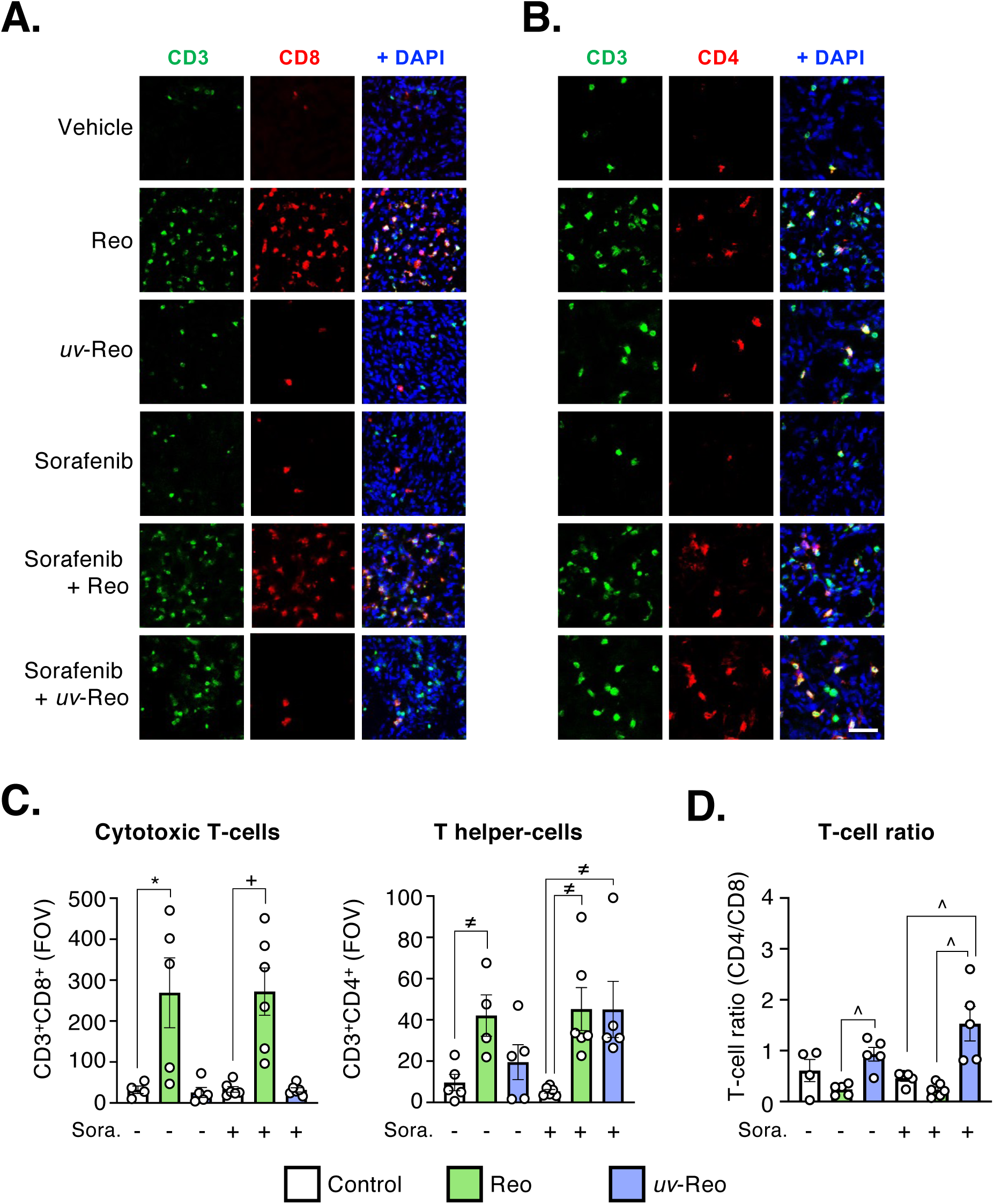
CD4^+^ T-helper cells, but not CD8^+^ CTLs, accumulate in murine hepatocellular carcinomas responding to the combination of *uv*-reovirus and sorafenib. (**A** – **B**) Representative images of 1MEA tumour cryosections taken from tumours harvested midway through the “therapy” phase, stained for CD3 and CD8 (*left*) or CD4 (*right*) then counterstained with DAPI. (**C**) Quantification of CD3^+^CD8^+^ (*left*) and CD3^+^CD4^+^ (*centre*) T-cells in random fields of view (FOV) taken from tumour cryosections in the indicated groups. (**D**) Comparison of CD4^+^:CD8^+^ T-cell ratio from quantification in ‘C’ (Scale bar = 50 µm; n = 5 mice per condition).

Cytokine array analysis (**Fig. S3A**) revealed increased intra-tumoural IFNG and TNFA (**Fig. 3A**) coincident with the accumulation of CD3^+^CD4^+^ T-cells. These cytokines are known to be expressed by T_H_1-activated T-cells and, accordingly, were detected in tumour-infiltrating CD3^+^CD4^+^ cells in mice treated with *uv*-Reo/sorafenib therapy (**Fig. 3B**). By contrast, levels of cytokines associated with T_H_2 and T_H_17 T-cell subsets did not mirror the pattern of CD4^+^ T-cell recruitment (**Fig. S3B**).

**Figure 3.**
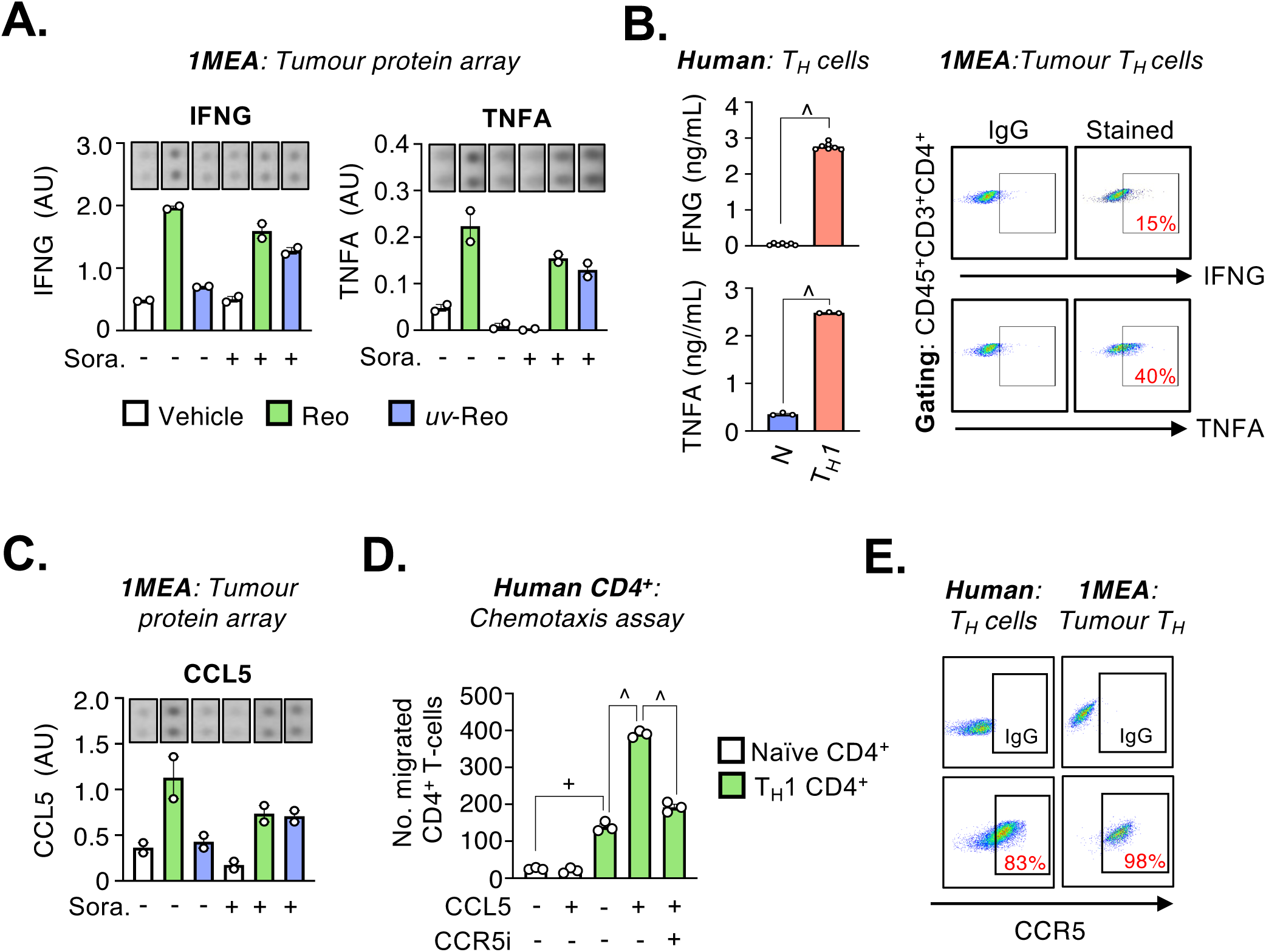
Treatment of murine HCC tumours with *uv*-Reo/sorafenib therapy elicits at CD4^+^ T_H_1-cell response. (**A**) Cytokine array data for the T_H_1-cytokines IFNG (*left*) and TNFA (*right*) from pooled tumour protein samples and from the indicated treatment groups. (**B**) ELISA-based quantification of IFNG and TNFA in supernatants from human CD4^+^ T-cells, *in vitro* (*left*), and flow cytometric detection of intra-cellular IFNG and TNFA in tumour-infiltration CD4^+^ T-cells in mice undergoing treatment with *uv*-Reo/sorafenib therapy (*right*). (**C**) Cytokine array data as described in ‘A’ but for CCL5. (**D**) Chemotaxis assay comparing the migratory potential of human CD4^+^ T-cells towards CCL5 and the dependency on CCR5, using the antagonist Maraviroc (CCR5i – 1µM) or vehicle (DMSO). (**E**) Flow cytometric detection of cell surface CCR5 on human T_H_1-activated CD4^+^ T-cells, *in vitro* (*left*), and tumour-infiltrating CD4^+^ T-cells in mice undergoing treatment with *uv*-Reo/sorafenib therapy (*right* – gated on CD45^+^CD3^+^CD4^+^ cells).

We also observed elevated levels of the T_H_1-cell chemokine CCL5 coincident with the increased abundance of CD3^+^CD4^+^ T-cells (**Fig. 3C**). *In vitro*, the expression of CCL5 by human and murine HCC cells was significantly increased by treatment with *uv*-Reo compared to Reo, even in the presence of sorafenib (**Fig. S3C - D**). CCL5 exerts a potent chemotactic effect on T_H_1-cells, but not naïve CD4^+^ T-cells, and a specific small molecule inhibitor supported that this chemotaxis was mediated by the chemokine receptor CCR5 (**Fig. 3D**). This was readily detected on the surface of T_H_1-activated T-cells *in vitro,* and on intra-tumoural CD3^+^CD4^+^ T-cells from mice treated with *uv*-Reo/sorafenib (**Fig. 3E**). These data indicate that the combination of *uv*-Reo and sorafenib exerts a suppressive effect on continuing HCC growth by skewing the intra-tumoural T-cell ratio in favour of CD4^+^ T_H_1-cells, not CD8^+^ CTLs.

### T_H_1-activated CD4^+^ T-cells exert contact-independent tumouricidal activity against HCC cells via a TNFA-dependent mechanism

We co-cultured HCC cells with CD4^+^ T-cells to determine whether the latter possessed anti-tumour properties. T_H_1-cells displayed measurable tumouricidal activity (**Fig. 4A**) that did not require the two cell types to be in physical proximity, demonstrating that a soluble factor was responsible (**Fig. 4B**). Thus, we examined the tumouricidal activity of soluble mediators released by T_H_1-cells known to have cytotoxic properties. We discovered that TNFA (**Fig. 4C – D**), not IFNG (**Fig. S4A**), was a direct-acting tumouricidal factor produced by T_H_1-cells. However, IFNG enhanced the sensitivity of HCC cells to TNFA killing by T_H_1-cells in co-culture assays (**Fig. S4B, Fig. 4E** – *left panel*).

**Figure 4.**
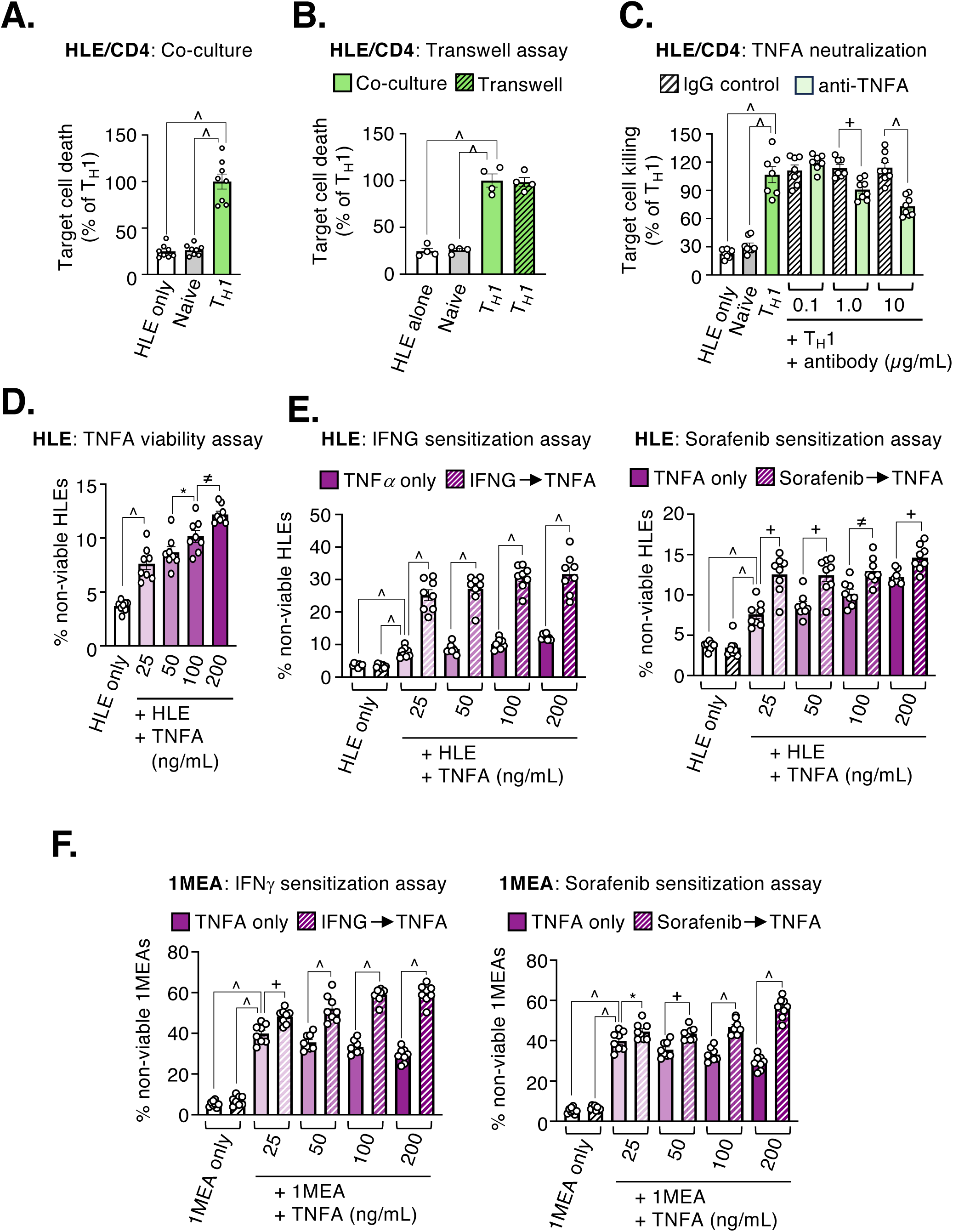
T_H_1-activated CD4^+^ T-cells exert a TNFA-dependent tumouricidal activity that is enhanced by tumour cell exposure to IFNG or sorafenib. (**A**) Flow cytometric quantification of target HLE killing by human CD4^+^ T-cells in direct co-culture or (**B**) following separation of T-cells/target cells by porous (0.4 µm) tissue culture inserts. (**C**) Quantification of HLE target killing in direct co-culture with CD4^+^ T-cells in the presence of neutralising antibodies against TNFA or IgG control. (**D**) Quantification of HLE cell viability in the presence of increasing concentrations of TNFA alone or (**E**) following pre-treatment with IFNG (*left* - 100 U/mL) or sorafenib (*right* - 7 µM). (**F**) Quantification of 1MEA cell viability in response to TNFA alone or following pre-treatment with IFNG (*left* – 100 U/mL) or sorafenib (*right* – 7 µM) (n = 4 – 8 per condition).

Next, we treated HCC cells with sorafenib, using a clinically relevant dose, to determine whether this could further modify the responses to TNFA and IFNG. Sorafenib significantly enhanced the anti-tumour activity of TNFA against human HCC cells (**Fig. 4E** – *right panel*), but not IFNG (**Fig. 4F** and **Fig. S4D**), and these findings were replicated using the mouse HCC cell line used for *in vivo*. A contribution from T_H_1-derived factors TRAIL and LTA was excluded based on antibody neutralisation experiments in co-culture assays (**Fig S4E – F**).

Together, these data demonstrated that TNFA was the dominant soluble mediator of T_H_1-cellular cytotoxicity towards HCC, with a tumour-sensitising role for both IFNG and sorafenib. However, because this difference alone did not explain the improved efficacy of the *uv*-Reo/Sorafenib combination, we reasoned that additional mechanisms likely enhance tumour killing for this treatment combination.

### IFNB induces a close-contact MHCII-independent tumouricidal activity in T_H_1-cells mediated by granzyme B/perforin

Type-I interferons are important mediators of anti-viral immunity *via* upregulation of interferon-stimulated genes (ISGs), as well as enhancing immune-mediated killing of infected cells. We, and others, have shown that Reo and *uv*-Reo are potent inducers of the type-I interferon, ‘IFNB’^7, 21^ particularly in the context of both HCC and primary liver tissue^21^. Thus, we compared the levels of IFNB produced by both human and mouse HCC cells *in vitro,* and mouse tumours *ex vivo,* when treated with Reo/*uv*-Reo monotherapy or in combination with sorafenib. Human and mouse HCC cells responded to *uv*-Reo (alone and in combination with sorafenib) with a robust induction of IFNB expression, which, surprisingly, was significantly larger than the response to Reo (**Fig. 5A - B**). Next, we examined the intracellular signalling events in HCC cells treated with Reo/*uv*-Reo alone and in the presence of sorafenib to better understand the differential induction of IFNB. The transcription factors NF*k*B p65 (RELA) and IRF3 were phosphorylated to a greater extent in HCCs treated with *uv*-Reo compared to Reo, both in the presence and absence of sorafenib, as determined by Western blot (**Fig. 5C**). Therefore, we compared the contribution made by each of these factors to the expression of IFNB using luciferase reporter assays containing either IRF3 (PD116) or NF*k*B p65 (PRDII) binding elements from the IFNB promoter. The IRF3 reporter was more strongly activated in HCC cells infected with *uv*-Reo (in both the presence and absence of sorafenib) compared to Reo (**Fig. S5A**). However, the opposite was true for the NF*k*B reporter with more robust activation in the presence of Reo, not *uv*-Reo (**Fig. S5B**). These data indicated that the differential activation and functional output from these pathways contributed the difference in IFNB induction by Reo/*uv*-Reo.

**Figure 5.**
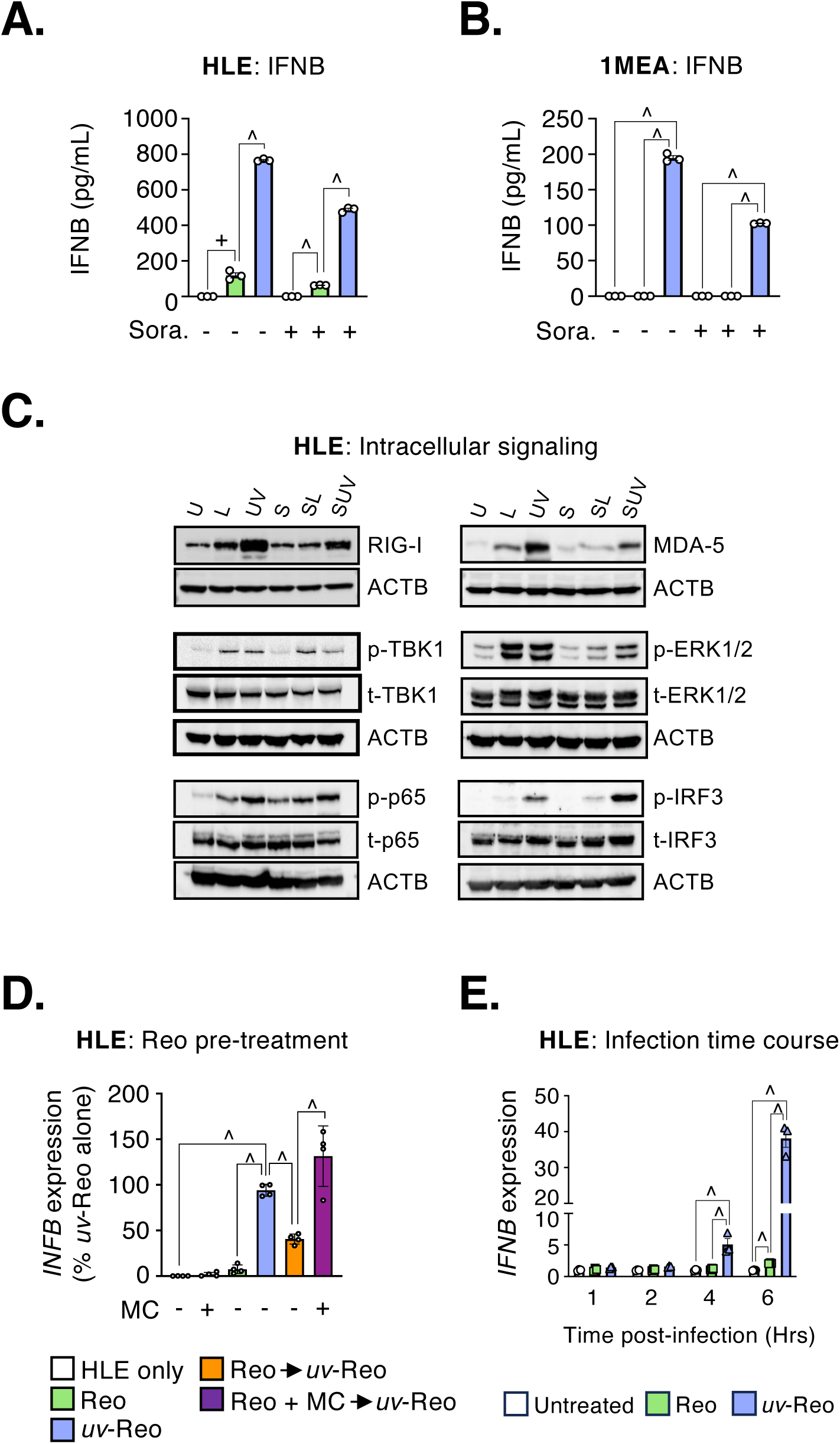
The differential induction of IFNB by Reo/*uv*-Reo can be attributed to virulence factor(s). Quantification of IFNB in supernatants from (**A**) human HLE cells or (**B**) mouse 1MEA cells, by ELISA. (**C**) Representative Western blots revealing activation of intracellular signalling pathways in HLEs following infection with Reo/*uv*-Reo alone or in combination with sorafenib, 24 hours post-infection (**D**) *IFNB* expression in HLE cells treated with Reo or *uv*-Reo, alone or sequentially, in the presence or absence of 2’-C-Methylcytidine. (**E**) Time-course of *IFNB* upregulation in HLE cells treated with Reo or *uv*-Reo (n = 3 per condition).

Viruses dedicate considerable proportions of their coding capacity towards the production of interferon antagonists within infected cells. Reovirus is no exception^22, 23^ so we hypothesised this may account for the differential induction of IFNB observed in HCC cells. Consistently, infecting HCC cells with Reo prior to *uv*-Reo significantly dampened the induction of IFNB compared to *uv*-Reo alone (**Fig. 5D**). In addition, the kinetics of virus-induced IFNB expression were slower with Reo compared to *uv*-Reo, indicating a delay in viral sensing (**Fig. 5E**). Critically, this effect was dependent upon Reo replication, and so presumably gene expression, as a nucleotide analogue inhibitor of the RNA-dependent RNA polymerase, 2’-C-methylcytidine, ameliorated the suppressive effect (**Fig. 5D**). These data indicated that a virulence factor could account for the differential induction of IFNB between Reo and *uv*-Reo, *in vitro*. Accordingly, Reo µNS protein is known to redirect IRF3 to viral replication factories^22^, consistent with decreased levels of phosphorylated protein within Reo infected HCC cells (**Fig. S5A**). Thus, whilst attenuation of the Type 3 Dearing strain may be severe compared to other *Orthoreoviruses*, it is by no means absent.

Interestingly, the highest level of intra-tumoural IFNB was found in tumours from mice treated with *uv*-Reo/sorafenib therapy (**Fig. 6A**), affirming that the magnitude of the IFNB response might play an important mechanistic role in suppressing tumour growth. Thus, we added recombinant IFNB to co-culture assays comprising HCC cells and T_H_1-cells to determine how this affected target cell killing. IFNB significantly enhanced the tumouricidal activity of T_H_1-cells (**Fig. 6B**) but, unlike TNFA, did so *via* a mechanism that required proximity between the two cell types (**Fig. 6C - D**). Next, we examined the effect of pre-treating HCC target cells with IFNB to determine whether the enhanced killing was a consequence of either increased target cell sensitivity or the augmented killing capacity of T_H_1-cells. Consistent with the latter, target cell killing was only enhanced when IFNB was added concurrently with T_H_1-cells (**Fig. 6E**) suggesting that the IFNB acted either directly on CD4^+^ T-cells alone or simultaneously on both cell types. Moreover, shRNA-mediated knockdown of IFNB in mouse 1MEA cells eliminated the efficacy of *uv*-Reo/Sorafenib therapy, *in vivo* (**Fig. 6F**) and this correlated with a failure to recruit CD4^+^ T-cells (**Fig. 6G**).

**Figure 6.**
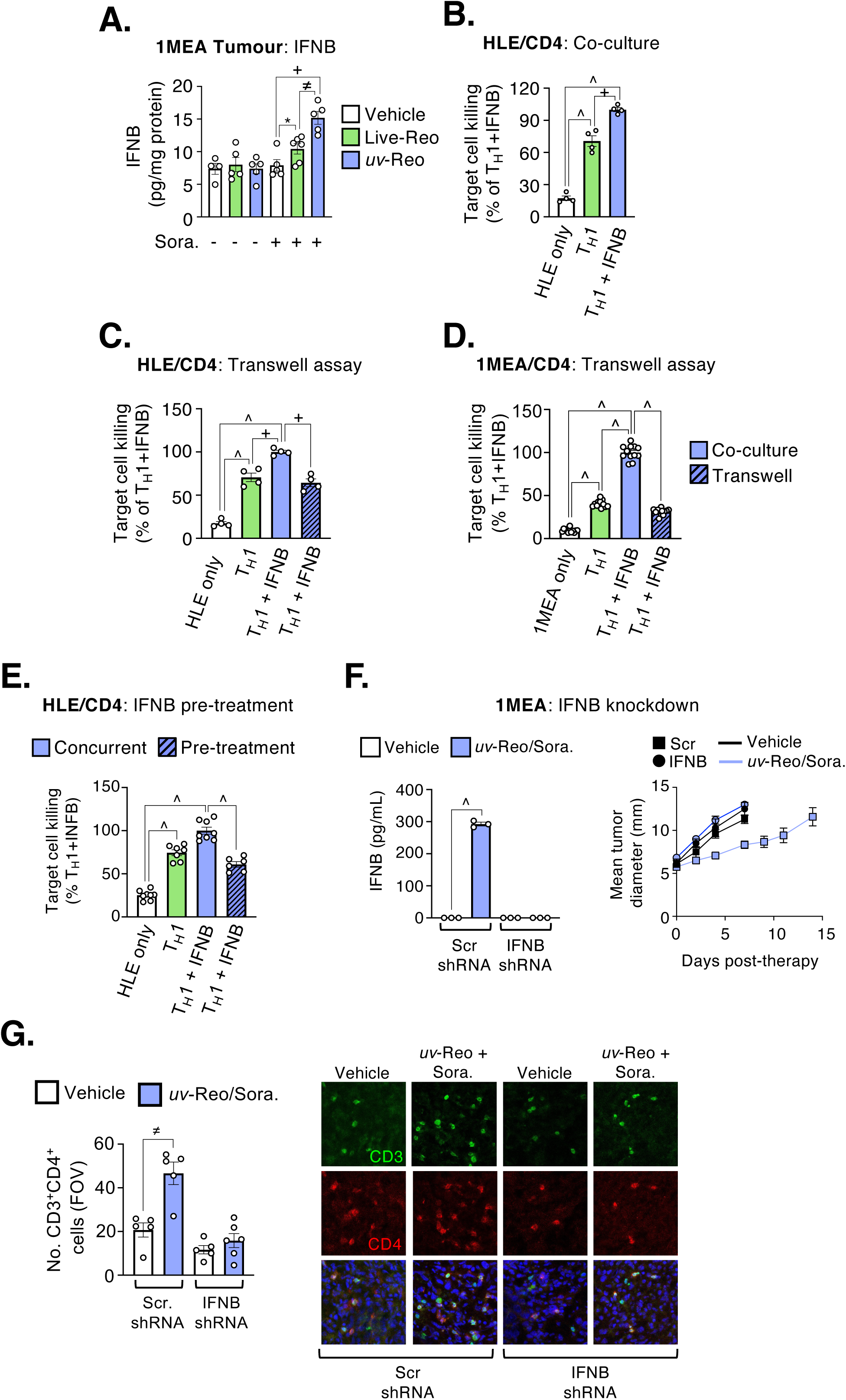
IFNB induces a proximity-dependent mode of tumour cell killing by T_H_1-cells and delays tumour growth, *in vivo*. (**A**) Quantification of IFNB in protein lysates from tumour-bearing mice treated with Reo/*uv*-Reo (2 pfu/cell) alone or in combination with sorafenib (7 µM). (**B**) Quantification of IFNB-induced HLE target cell killing by T_H_1-activated CD4^+^ T-cells in direct co-culture or following separation of T-cells/target cells by porous tissue culture inserts in (**C**) human and (**D**) mouse systems. (**E**) Quantification of IFNB-induced killing of HLE target cells when co-administered with T-cells or given to target cells as a pre-treatment. (n = 4 – 8 per condition). (**F**) Quantification of IFNB released by 1MEA cells carrying Scrambled or IFNB-targeted shRNAs following treatment with *uv*-Reo (*left*) and their response to *uv*-Reo/Sorafenib therapy, *in vivo* (*right*). (**G**) Quantification (*left*) and representative images (*right*) of CD3^+^CD4^+^ T-cell abundance in 1MEA tumours carrying Scrambled or IFNB-targeted shRNA constructs treated with *uv*-Reo/Sorafenib therapy.

We next investigated potential juxtacrine mediators of IFNB-enhanced T_H_1-mediated killing of HCC cells, namely TNF-related apoptosis-inducing ligand (TRAIL) and FasL/Fas. However, inhibiting these pathways in co-culture assays did not inhibit target cell killing (**Fig. S6A – G**). Consequently, we considered the involvement of MHCII-dependent degranulation, described previously for cytotoxic CD4^+^ T-cells^24^. However, MHCII was absent on both human and mouse HCC targets in culture, and addition of MHCII-neutralising antibody to co-culture assays had no effect (**Fig. 7A - B**). Despite the lack of MHCII involvement, addition of EGTA (a calcium chelating inhibitor of degranulation) to co-culture assays dose-dependently inhibited IFNB-enhanced target cell killing (**Fig. 7C**). Hence, we next quantified the proportion of CD4^+^ T-cells with surface expression of the degranulation marker CD107a and intra-cellular expression of both granzyme B and perforin, important components of the apoptosis-inducing machinery deployed by cytotoxic T-cells. We observed a marked increase in the proportion of human T-cells with surface CD107a expression and intra-cellular granzyme B (GZMB) following T_H_1-activation and this was further enhanced upon stimulation with IFNB (**Fig. 7D – E**). A smaller, but statistically significant, change in intra-cellular perforin (PRF) was also detected. Similar results were obtained for mouse CD4^+^ T-cells (**Fig. S7A**). GZMB and PRF were readily detected in supernatants from T_H_1-activated T-cells, and levels were further increased upon treatment with IFNB (**Fig. 7F**). Accordingly, addition of a small molecule GZMB inhibitor to co-culture assays dose dependently inhibited IFNB-enhanced killing (**Fig. 7G** – *left panel*). The same response was also seen in the presence of a pan-caspase inhibitor (**Fig. 7G** – *right panel*). Taken together, these data demonstrate that IFNB enhanced the tumouricidal activity of CD4^+^ T-cells by increasing the expression and/or licensing of GZMB/PRF activity against MHCII-negative HCC cells.

**Figure 7.**
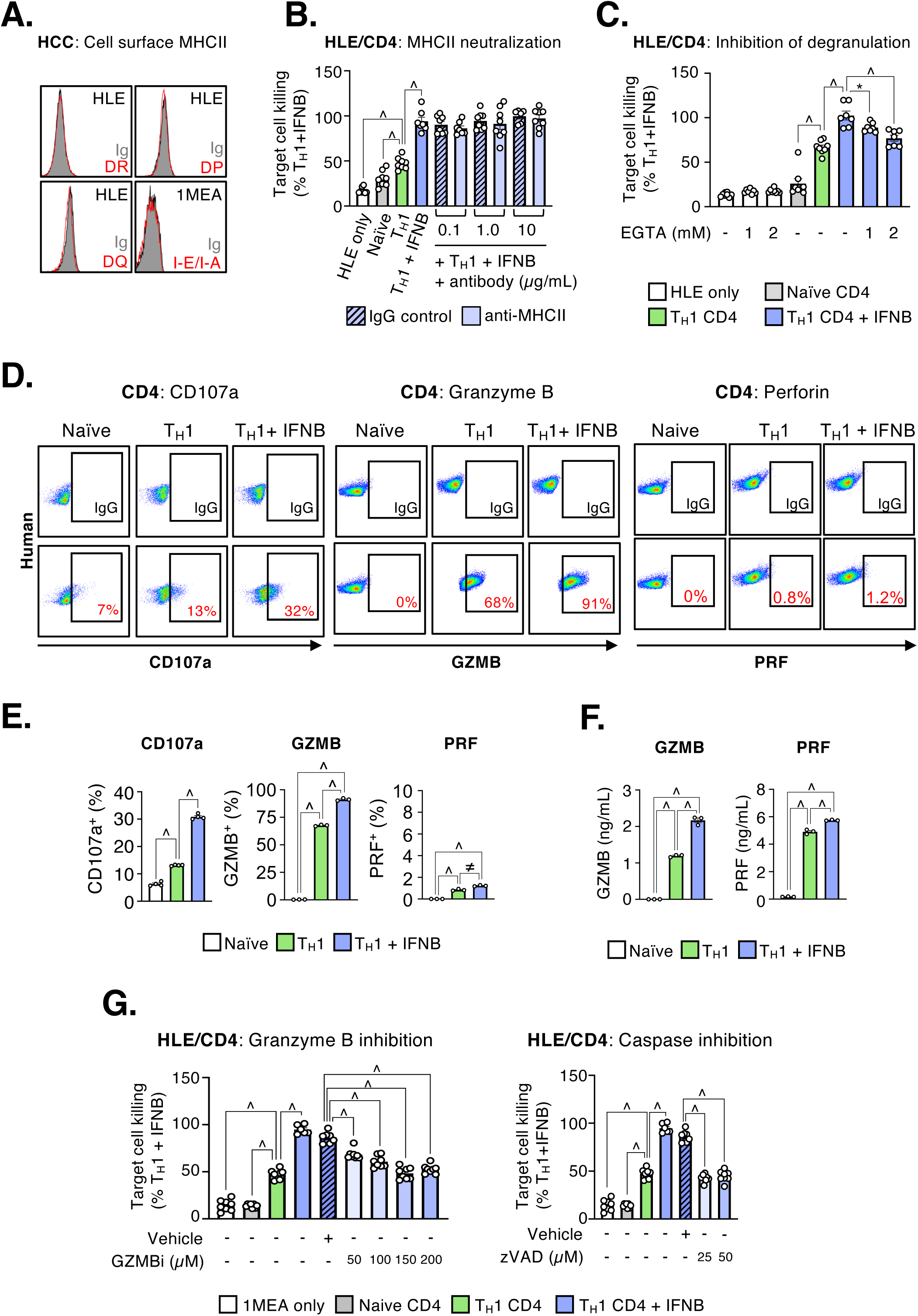
IFNB induces an antigen and perforin-independent, granzyme B-dependent mode of tumouricidal activity in T_H_1-activated CD4^+^ T-cells. (**A**) Flow cytometric detection of cell surface MHCII proteins on human and murine HCC cells, *in vitro*. (**B**) Quantification of HLE target cell killing by human T_H_1-activated CD4^+^ T-cells induced by IFNB in the presence of neutralising antibodies against MHCII or (**C**) the degranulation inhibitor EGTA. (**D**) Representative flow cytometry plots showing detection of cell surface CD107a (*left*) and intra-cellular GZMB (*centre*) or PRF (*right*) in human CD4^+^ T-cells alongside (**E**) quantification of the proportion cells positive for each marker. (**F**) Quantification of cell-free GZMB and PRF in supernatants from human CD4^+^ T-cells, by ELISA. (**G**) Quantification of HLE target cell killing by human T_H_1-activated CD4^+^ T-cells in the presence of IFNB and a GZMB inhibitor - z-AAD-CMK (*left*) or caspase inhibitor - z-VAD-FMK (*right*). (n = 4 – 8 per condition).

Finally, given that CD8^+^ cells were also present, but not enriched, within tumours treated with *uv*-Reo and sorafenib (**Fig. 2**), we compared their ability to engage in antigen-independent HCC killing with that of CD4^+^ T-cells, *in vitro*. Consistent with the notion that CD8^+^ T-cells played a lesser role in tumour response to therapy, whilst they could indeed kill HCCs upon activation and this was further enhanced by IFNB, their tumouricidal activity was significantly lower than that of CD4^+^ T-cells (**Fig. S7B**). Furthermore, CD8^+^ T-cell tumouricidal activity required neither TNFA (**Fig. S7C**) nor degranulation (**Fig. S7D**) and was therefore distinct mechanistically from that of CD4^+^ T-cells.

Thus, we conclude that the magnitude of the initial IFNB response, resulting from a lack of viral antagonist, underpins the improved efficacy of the *uv*-Reo/Sorafenib therapy in preclinical HCC models. The induction of IFNB was associated with increased expression of CCL5 and supported the accumulation of T_H_1-activated CD4^+^ T-cells with basal tumouricidal activity, facilitated by their expression of TNFA and IFNG. Crucially, IFNB licenses and/or focusses the cytolytic activity of GZMB and PRF, derived from T_H_1-cells, against MHCII-negative tumours.

## DISCUSSION

This study demonstrates the superior therapeutic efficacy of a unique immunotherapy combining *uv*-inactivated reovirus with the targeted agent sorafenib over either sorafenib monotherapy or combination with live Reo, as a treatment for HCC. This combination therapy significantly extended the survival of mice bearing HCC tumours by engaging a multi-faceted, MHCII-independent response from cytotoxic CD4^+^ T_H_-cells (CTHs), induced by IFNB.

The *uv*-Reo/sorafenib therapy specifically induced increased expression of the chemokine CCL5 and cytokine IFNB relative to other treatment modalities. This was accompanied by increased abundance of intra-tumoural CCR5^+^ T_H_1-activated CD4^+^ T-cells, skewing the T-cell ratio in favour of T_H_1-cells. Tumour-infiltrating lymphocytes (TILs) exerted basal levels of paracrine tumouricidal activity *via* TNFA, which was enhanced in HCC cells by both IFNG and sorafenib. Concomitantly, increased expression of IFNB in HCC cells following stimulation with *uv*-Reo/sorafenib was a crucial mechanistic switch, stimulating MHCII-independent degranulation involving GZMB and PRF, requiring close cell-cell contact. Hence, our findings support that *uv*-inactivated reovirus could be used to enhance the efficacy of sorafenib during the treatment of HCC by engaging MHCII-independent GZMB^+^PRF^+^ T_H_1-cells.

The role of TNF in liver cancer is complex and studies can be contradictory. A considerable body of evidence details a pro-tumourigenic role for TNF in the development of liver cancer^25^. However, in pre-clinical cancer models, direct intra-tumoural injection of TNF resulted in widespread necrosis, an effect attributed to its anti-vascular activity, rather than direct tumour cytolysis^26–28^. The consensus is, therefore, that TNF is a weak direct-acting cytolytic, consistent with findings herein. However, this limited tumouricidal activity can be significantly enhanced by co-administration of IFNG, a phenomenon demonstrated in a variety of tumour types, including melanoma, breast, and colon cancer^26, 27^, confirmed again here. We now add that sorafenib also sensitised HCCs to TNFA-induced cell death, reminiscent of responses described previously for other TNF family members including TRAIL and Fas, attributed to downregulation of the anti-apoptotic protein, Mcl-1^29^ in tumour cells. Although it is likely that the response to TNFA is enhanced by sorafenib in a similar way, the precise mechanism has yet to be determined. Interestingly, we found that TNFA/IFNG-mediated HCC killing by T_H_1-cells occurred in the presence of the pan-caspase inhibitor Z-VAD-FMK, implicating a caspase-independent pathway.

These data provide an important mechanistic insight and suggest that the efficacy of *uv*-Reo/sorafenib therapy is mediated, at least in part, by a skewing of T-cell ratios in favour of IFNG^+^ TNFA^+^ T_H_1-cells. However, IFNG and TNFA alone do not fully explain how T_H_1-cells control tumour growth in response to *uv*-Reo/sorafenib and not in other treatment groups where they are also equally abundant.

We, and others, have demonstrated that *uv*-Reo elicits a significantly more robust induction of IFNB in treated cells compared with Reo^21, 30^, confirmed again, here, in both human and murine HCC cells. Our analysis indicates a Reo-derived antagonist virulence factor is responsible for the differential induction of IFNB by Reo/*uv*-Reo, as well as the ability of Reo to suppress ensuing responses (**Fig 5C**). Several factors have been reported including the non-structural mu protein (*µ*NS)^22^ and the outer capsid *σ*3^23^. It may be that one or both contribute to suppression of IFNB in HCCs infected by Reo, although increased IRF3 phosphorylation in cells exposed to *uv*-Reo is consistent with a lack of *µ*NS function, rather than *σ*3 which predominantly antagonises PKR.

The genetic determinants of reovirus strain variability, specifically relating to interferon antagonism and apoptotic potential, have been identified^31, 32^. Type-3 Dearing strain (T3D) shows a reduced ability to suppress type-I interferon responses^31, 32^ and demonstrates enhanced pro-apoptotic activity in infected cells^31^. This strain has been extensively evaluated as a oncolytic agent in clinical trials. However, in the context of HCC, T3D’s attenuation appears insufficient, failing to elicit a robust interferon response or effectively control tumour growth. This finding suggests that many natural or genetically attenuated OVs currently in clinical trials might not be adequately attenuated for all cancer types, particularly those that are considered immunologically cold. This raises the question of whether the enhanced IFNB response induced by *uv*-inactivated viruses could lead to superior clinical efficacy in these cancer types. Interestingly, the TKI sorafenib was key to the robust induction of IFNB that underpinned the CD4^+^ T-cell response, despite being naturally immunosuppressive. This raises the question of whether *uv*-inactivated viruses might be effective in other cancer types treated with sorafenib and similar TKIs.

This study demonstrates that IFNB induces a unique GZMB/PRF-mediated tumouricidal activity in T_H_1-cells, independent of MHCII expression by target HCCs. Although the expression of GZMB and PRF by CD4^+^ T-cells is reported in the context of viral infection, this mode of killing is both antigen-specific and reliant on MHCII^33, 34^. How an immune synapse is able to form between IFNB-stimulated T_H_1-cells and MHCII-negative tumour cells, thereby inducing target cell death, is not fully understood but, a mechanism involving the upregulation of such stress-induced NKG2D (present upon activated T_H_1-cells) ligands, in conjunction with ICAM1/LFA1, has been proposed^35–39^. Interestingly, NKG2D ligands can by induced within tumour cells following viral infection and in response to type-I interferon^38, 40^.

CD8^+^ T-cells were also present in tumours treated with *uv*-Reo/sorafenib therapy, although their abundance was unchanged relative to controls. Furthermore, CD8^+^ T-cells were surprisingly less tumouricidal than their CD4^+^ counterparts in the context of antigen-independent immunity, and utilised neither TNFA nor GZMB. These data indicate that the immediate response to therapy with *uv*-Reo/sorafenib is dominated by T_H_1-activated T-cells. Despite this, we do not discount a major role for antigen-specific immunity during the post-therapy phase, which contributed significantly to the overall survival advantage (Fig 1). How initial MHCII-independent responses translate into adaptive immunity remains the focus of ongoing research.

Taken together, our data demonstrate that *uv*-inactivated reovirus and the targeted agent, sorafenib, combine to drive IFNB and CCL5 mediated tumour infiltration by MHCII-independent GZMB^+^PRF1^+^ T_H_1-cells, capable of establishing subsequent long-lasting tumour survival despite the cessation of therapy. This also highlights an overlooked role for CD4^+^ T_H_1-cells in mediating MHC-independent anti-tumour immunity. Further mechanistic elaboration is needed to fully understand how non-canonical effector cells can be further exploited as a therapeutic tool for the treatment of HCC.

## Supporting information

All supplemental information

## ACKNOWLEDGEMENTS

*Funding*: The authors thank the Medical Research Council (Award No: MR/T016205, awarded to SG), and Cancer Research UK (Award No: 29039, awarded to AS) for funding this research. We also thank the NHS Blood and Transplant Service for the provision of donor blood samples, the St. James’s Biological Services Unit (SBS) for their support with animal welfare and husbandry, and Oncolytics Biotech Inc. for supplying clinical grade human *Orthoreovirus*, type-3 Dearing strain, Pelareorep.

## Notes

**COMPETING INTERESTS** All authors declare that there are no conflicts of interest.

### Competing Interest Statement

The authors have declared no competing interest.

### Summary of Updates

This version provides additional evidence of the importance of IFNB, whereby tumours lacking expression are unresponsive to therapy. Moreover, we show that infiltrating cytotoxic CD4+ cells are integral to enhanced HCC survival, killing tumour cells via an IFNB-driven, MHCII independent juxtacrine mechanism using granzyme B and perforin.

