## Supplementary material for "IFNB induced by non-lytic virus immunotherapy promotes improved survival in hepatocellular carcinoma, mediated by MHCII-independent cytotoxic CD4^+^ T-cells": All supplemental information

Fig. S1

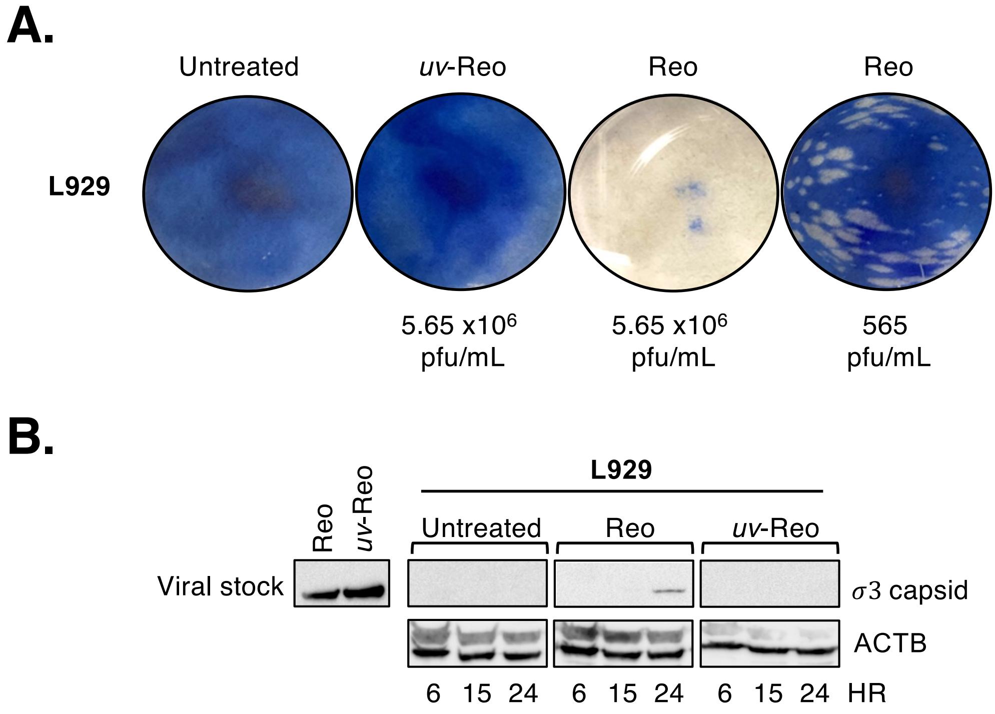
A.
Untreated
uv-Reo
Reo
Reo
L929
5.65 x10^6 pfu/mL
5.65 x10^6 pfu/mL
565 pfu/mL
B.
L929
Reo
uv-Reo
Untreated
Reo
uv-Reo
Viral stock
σ3 capsid
ACTB
6 15 24 6 15 24 6 15 24 HR

**Figure S1. *uv*-irradiation renders Reovirus replication deficient.** (**A**) Representative images of a plaque assay performed using Reo or *uv*-Reo on the permissive L929 mouse fibroblast cell line, images taken five days post-infection. (**B**) Western blot for $\sigma$3 capsid protein performed on lysates extracted from L929 cells at the indicated times post-infection with either Reo or *uv*-Reo, and on viral stocks.

### Figure S2

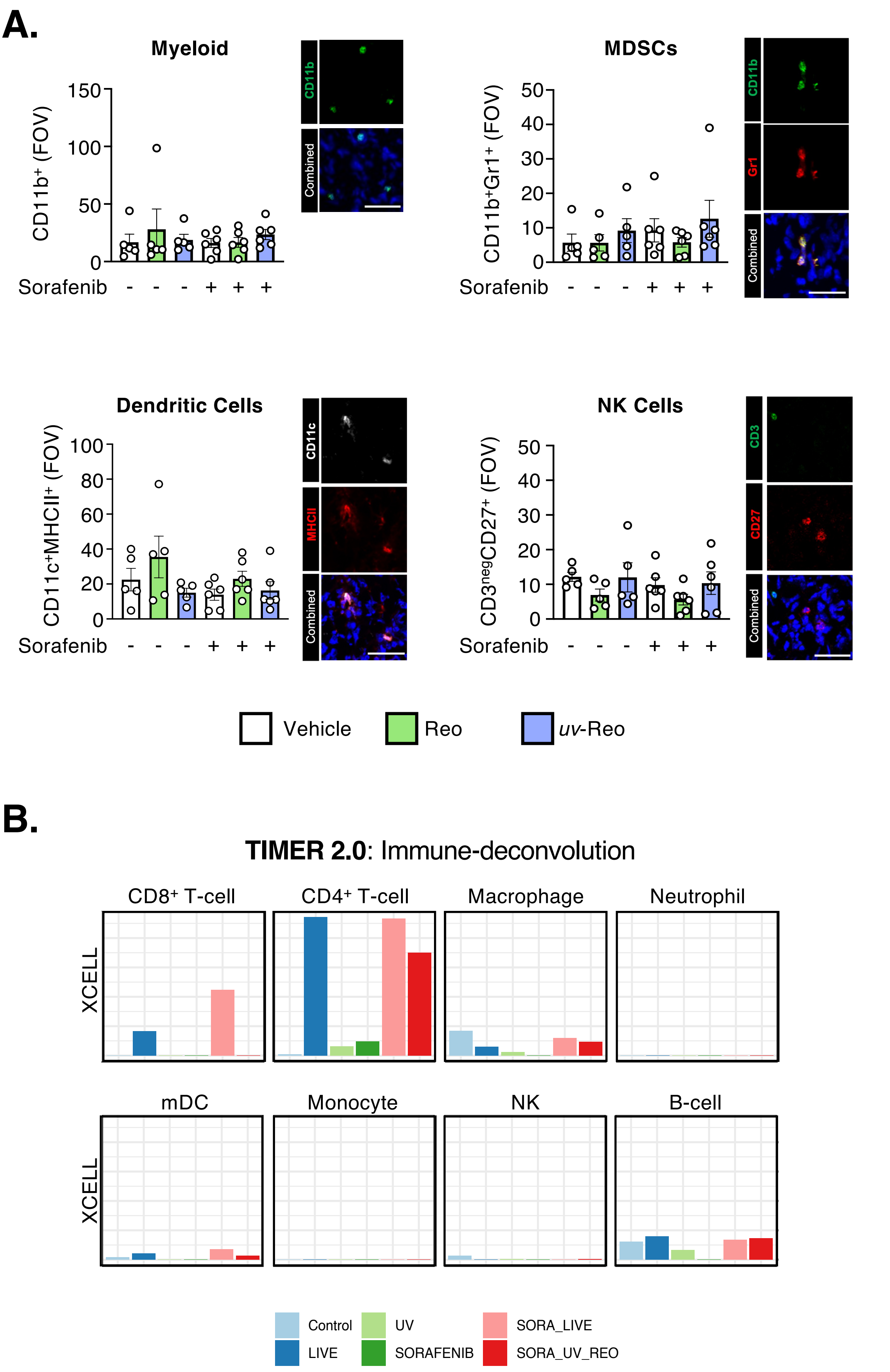
A.
Myeloid
CD11b⁺ (FOV)
150
100
50
0
Sorafenib - - - + + +
CD11b
Combined
MDSCs
CD11b⁺Gr1⁺ (FOV)
50
40
30
20
10
0
Sorafenib - - - + + +
CD11b
Gr1
Combined
Dendritic Cells
CD11c⁺MHCII⁺ (FOV)
100
80
60
40
20
0
Sorafenib - - - + + +
CD11c
MHCII
Combined
NK Cells
CD3negCD27⁺ (FOV)
50
40
30
20
10
0
Sorafenib - - - + + +
CD3
CD27
Combined
Vehicle
Reo
uv-Reo
B.
TIMER 2.0: Immune-deconvolution
CD8⁺ T-cell
CD4⁺ T-cell
Macrophage
Neutrophil
XCELL
mDC
Monocyte
NK
B-cell
XCELL
Control
LIVE
UV
SORAFENIB
SORA_LIVE
SORA_UV_REO

**Figure S2. Leukocyte abundance in syngeneic mouse hepatocellular carcinomas is largely unaffected by treatment with either reovirus or sorafenib.** (**A**) Cryosections were taken from murine HCC tumors grown in syngeneic hosts during the "Therapy-phase" during treatment with either Reo/*uv*-Reo alone ($1x10^7$ pfu) or in combination with sorafenib (7 $\mu$M) or vehicle. Tumor cryosections were immunolabelled for the detection of pan-myeloid cells ($CD11b^+$), MDSCs ($CD11b^+Gr1^+$), dendritic cells ($CD11c^+MHCII^+$) or NK cells ($CD3^{neg}CD27^+$) (**B**). RNASeq data obtained from 1MEA tumours in mice treated with Reo/*uv*-Reo alone or in combination with sorafenib was subjected to immune cell estimation using the XCELL algorithm, using the TIMER2.0 immune estimation function (n = 5 mice per condition).

#### Figure S3

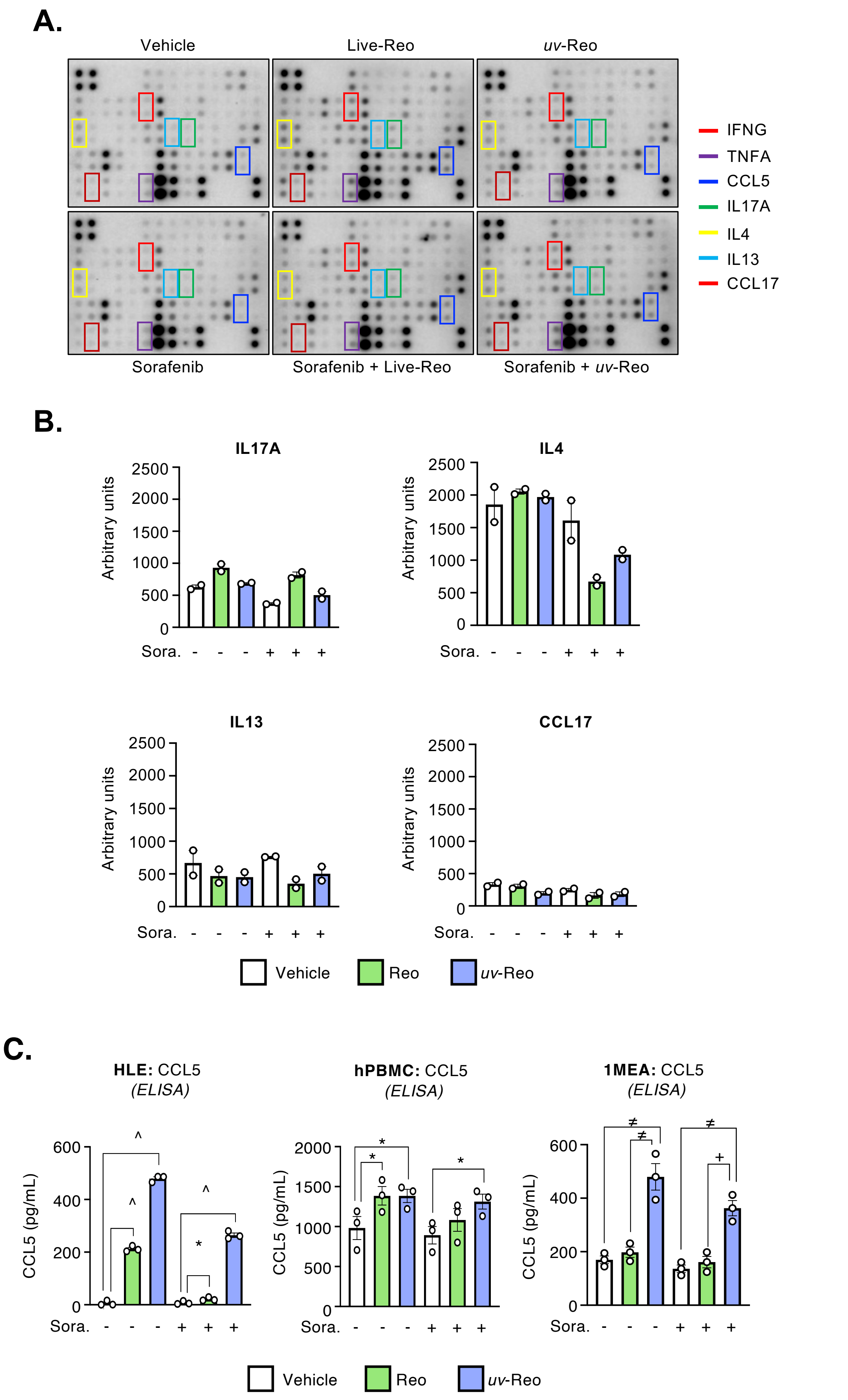

A.
Vehicle
Live-Reo
uv-Reo
Sorafenib
Sorafenib + Live-Reo
Sorafenib + uv-Reo
IFNG
TNFA
CCL5
IL17A
IL4
IL13
CCL17
B.
IL17A
IL4
IL13
CCL17
Arbitrary units
Sora. - - - + + +
Vehicle
Reo
uv-Reo
C.
HLE: CCL5 (ELISA)
hPBMC: CCL5 (ELISA)
1MEA: CCL5 (ELISA)
CCL5 (pg/mL)
Sora. - - - + + +
Vehicle
Reo
uv-Reo

**Figure S3. Reovirus alone or in combination with sorafenib induced an intra-tumoral cytokine profile inconsistent with the recruitment or either $T_H2$ or $T_H17$ $CD4^+$ T-cell.** (**A**) Images of cytokine arrays generated using pooled total tumor protein extracts from murine 1MEA HCC tumors grown in syngeneic host collected during the "Therapy-phase" following treatment with Reo/*uv*-Reo alone ($1x10^7$ pfu) or in combination with sorafenib (10 mg/Kg) or vehicle. Cytokines of interest are highlighted. (**B**) Semi-quantitative data for $T_H2$ and $T_H17$cytokines from protein arrays shown in 'A'. (Pooled lysates were generated from 4 – 5 mice per treatment group and each cytokine semi-quantitatively measured in duplicate) (**C**) CCL5 was quantified in supernatants from human HLE monolayer cells (*left*), human PBMCs (*centre*), and mouse 1MEA cells grown as spheroids (*right*) following overnight treatment with Reo/*uv*-Reo alone (2 pfu/cell) or in combination with sorafenib (7 $\mu$M) (* $p<0.05$, ≠ $p<0.01$, + $<0.001$, ^ $<0.0001$; n = 3 per condition).

### Figure S4

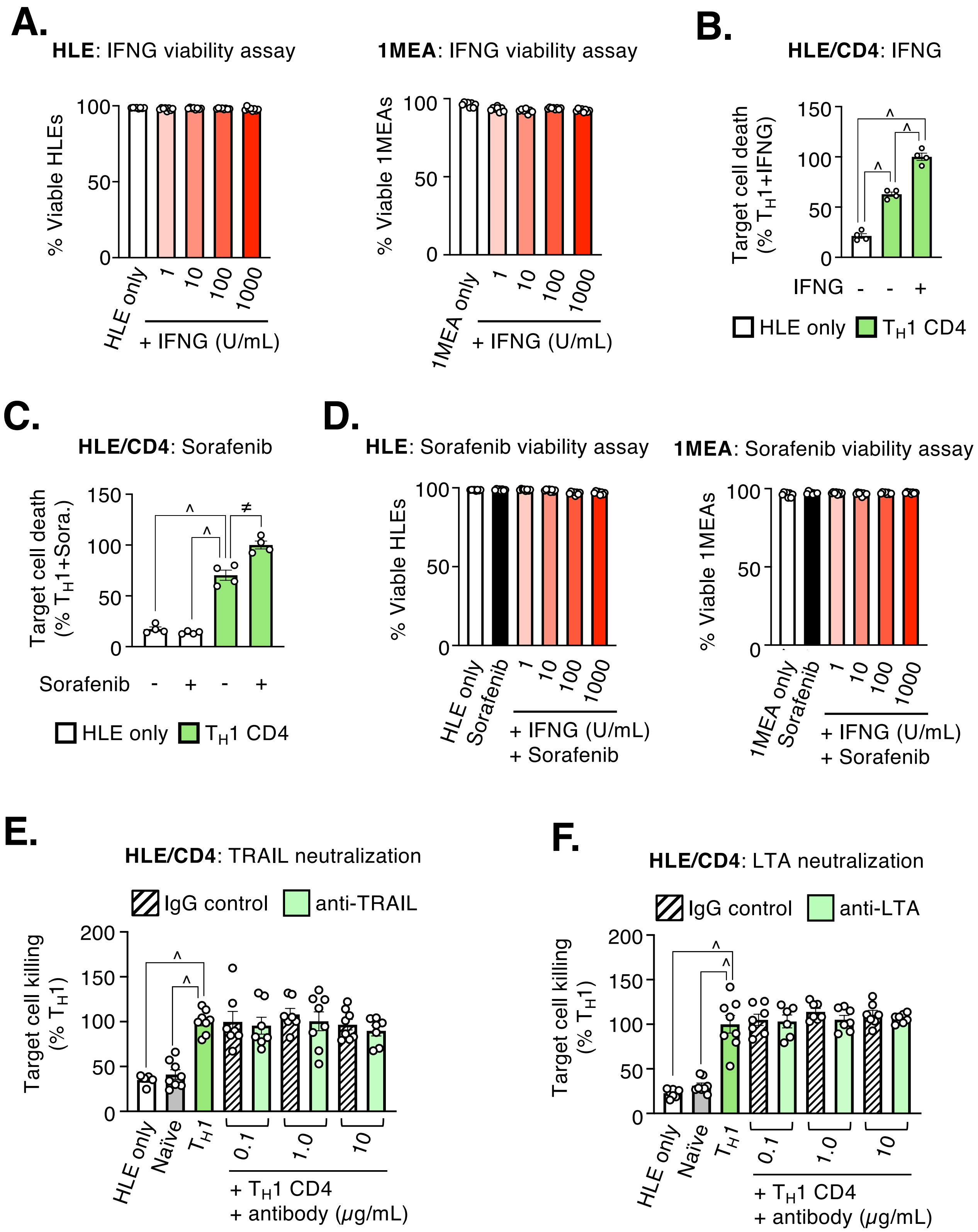
A.
HLE: IFNG viability assay
% Viable HLEs
HLE only
+ IFNG (U/mL)
1MEA: IFNG viability assay
% Viable 1MEAs
1MEA only
+ IFNG (U/mL)
B.
HLE/CD4: IFNG
Target cell death (% TH1+IFNG)
IFNG - - +
HLE only
TH1 CD4
C.
HLE/CD4: Sorafenib
Target cell death (% TH1+Sora.)
Sorafenib - + - +
HLE only
TH1 CD4
D.
HLE: Sorafenib viability assay
% Viable HLEs
HLE only
Sorafenib
+ IFNG (U/mL)
+ Sorafenib
1MEA: Sorafenib viability assay
% Viable 1MEAs
1MEA only
Sorafenib
+ IFNG (U/mL)
+ Sorafenib
E.
HLE/CD4: TRAIL neutralization
IgG control
anti-TRAIL
Target cell killing (% TH1)
HLE only
Naïve
TH1
0.1
1.0
10
+ TH1 CD4
+ antibody (μg/mL)
F.
HLE/CD4: LTA neutralization
IgG control
anti-LTA
Target cell killing (% TH1)
HLE only
Naïve
TH1
0.1
1.0
10
+ TH1 CD4
+ antibody (μg/mL)

**Figure S4. The tumoricidal activity of $T_H$1-cells against HCC cells is enhanced by pre-treatment with IFNG or sorafenib and killing is mediated by neither TRAIL nor LTA. (A)** Human and murine HCC cells were treated with increasing concentration of IFNG overnight and tumor cell viability determined by flow cytometry. (**B**) Co-culture assay between human HLE HCC cells pre-treated with or without IFNG (100 U/mL) and $T_H$1-activated $CD4^+$ T-cells. Target HCC cell viability determined by flow cytometry following overnight incubation (**C**) Co-culture assay between human HCC cells pre-treated with or without sorafenib (7 $\mu$M) and $T_H$1-activated $CD4^+$ T-cells. Target HCC cell viability was determined by flow cytometry following overnight incubation. (**D**) Human and murine HCC cells pre-treated with sorafenib (7 $\mu$M) prior to exposure to increasing concentrations of IFNG overnight. HCC viability was determined by flow cytometry. Co-culture assays were performed overnight between human HLE HCC cells and either naive or $T_H$1-activated $CD4^+$ T-cells in the presence of increasing concentrations of neutralizing antibodies against either TRAIL (**E**) or LTA (**F**) and target HCC cell viability was determined by flow cytometry (* $p<0.05$, ≠ $p<0.01$, + <0.001, ^ <0.0001; n = 4 – 8 per condition).

### Figure S5

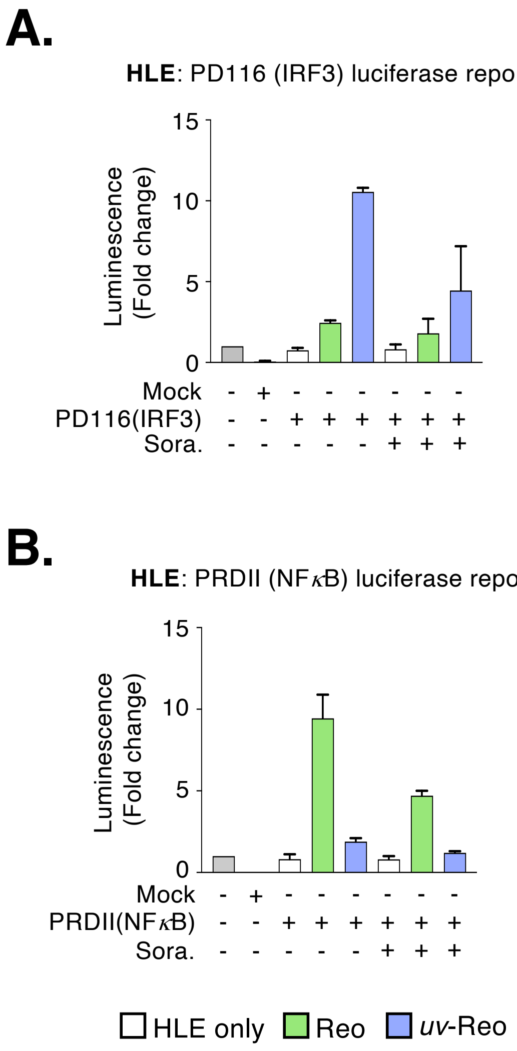
A.
HLE: PD116 (IRF3) luciferase reporter
Luminescence (Fold change)
15
10
5
0
Mock - + - - - - - -
PD116(IRF3) - - + + + + + +
Sora. - - - - - + + +
B.
HLE: PRDII (NFκB) luciferase reporter
Luminescence (Fold change)
15
10
5
0
Mock - + - - - - - -
PRDII(NFκB) - - + + + + + +
Sora. - - - - - + + +
HLE only
Reo
uv-Reo

**Figure S5. Differential activation of intracellular signalling in HCC following infection by Reo/*uv*-Reo.** HLE cells were transfected with luciferase reporter constructs containing (**A**) IRF3 and (**B**) NF*k*B responsive elements from the IFNB promoter and treated with Reo/uv-Reo (2 pfu/cell) alone or in combination with sorafenib (7 μM). (* $p<0.05$, ≠ $p<0.01$, + <0.001, ^ <0.0001; n = 2 - 3 per condition).

### Figure S6

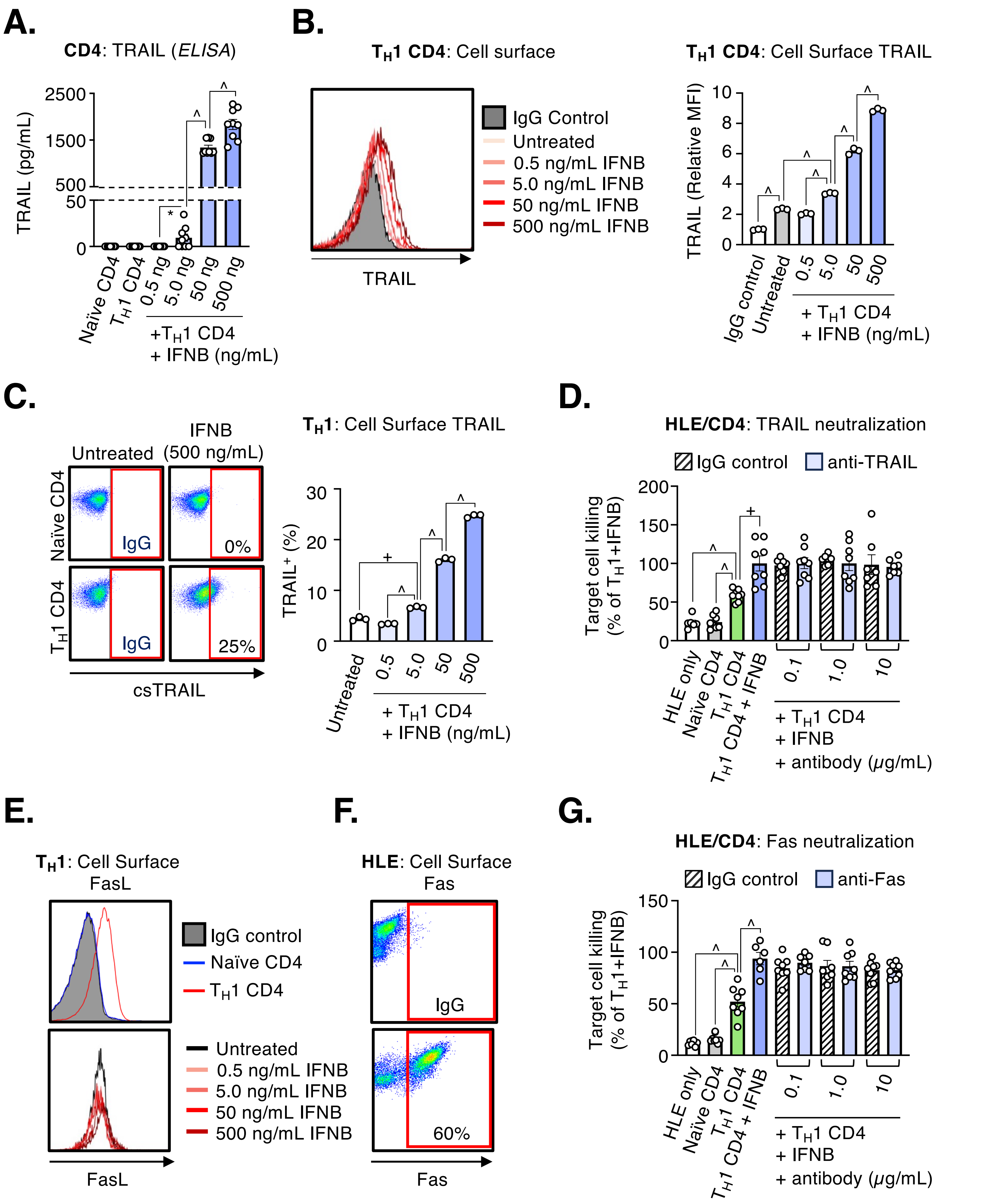
A.
CD4: TRAIL (ELISA)
TRAIL (pg/mL)
2500
1500
500
50
0
Naïve CD4
TH1 CD4
0.5 ng
5.0 ng
50 ng
500 ng
+TH1 CD4
+ IFNB (ng/mL)
B.
TH1 CD4: Cell surface
IgG Control
Untreated
0.5 ng/mL IFNB
5.0 ng/mL IFNB
50 ng/mL IFNB
500 ng/mL IFNB
TRAIL
TH1 CD4: Cell Surface TRAIL
TRAIL (Relative MFI)
IgG control
Untreated
+ TH1 CD4
+ IFNB (ng/mL)
C.
Untreated
IFNB (500 ng/mL)
Naïve CD4
TH1 CD4
IgG
0%
25%
csTRAIL
TH1: Cell Surface TRAIL
TRAIL+ (%)
D.
HLE/CD4: TRAIL neutralization
IgG control
anti-TRAIL
Target cell killing (% of TH1+IFNB)
HLE only
Naïve CD4
TH1 CD4
TH1 CD4 + IFNB
+ TH1 CD4
+ IFNB
+ antibody (μg/mL)
E.
TH1: Cell Surface FasL
IgG control
Naïve CD4
TH1 CD4
Untreated
0.5 ng/mL IFNB
5.0 ng/mL IFNB
50 ng/mL IFNB
500 ng/mL IFNB
FasL
F.
HLE: Cell Surface Fas
IgG
60%
Fas
G.
HLE/CD4: Fas neutralization
IgG control
anti-Fas
Target cell killing (% of TH1+IFNB)
HLE only
Naïve CD4
TH1 CD4
TH1 CD4 + IFNB
+ TH1 CD4
+ IFNB
+ antibody (μg/mL)

**Figure S6. The tumoricidal activity of $T_H$1-activated $CD4^+$ T-cells treated with IFNB is not mediated by TRAIL or FasL/Fas.** (**A**) $T_H$1-activated $CD4^+$ T-cells were treated with increasing concentrations of IFNB overnight and the concentration of soluble TRAIL was measured in supernatants by ELISA (n = 9). (**B - C**) $T_H$1-activated $CD4^+$ T-cells were treated overnight as described in 'A' and cell surface expression of TRAIL was quantified by flow cytometry (n = 3). (**D**) Co-culture assay between human HLE HCC cells and either naive or $T_H$1-activated $CD4^+$ T-cells with or without IFNB in the presence of increasing concentrations of anti-human TRAIL neutralizing antibody. Following incubation overnight, target HCC cell killing was determined by flow cytometry (n = 8). (**E**) Human HLE HCC cells were treated with increasing concentrations of IFNB overnight and cell surface expression of FasL was quantified by flow cytometry (n = 3). (**F**) Expression of FasL receptor – Fas – on human HLE HCC cells was confirmed by flow cytometry (n = 3). (**G**) Co-culture assay between human HLE HCC cells and either naive or $T_H$1-activated $CD4^+$ T-cells, with or without IFNB, in the presence of increasing concentrations of anti-human Fas neutralizing antibody or isotype matched control (csTRAIL = cell surface TRAIL; * $p<0.05$, ≠ $p<0.01$, + $<0.001$, ^ $<0.0001$; n = 8).

### Figure S7

A.

B.

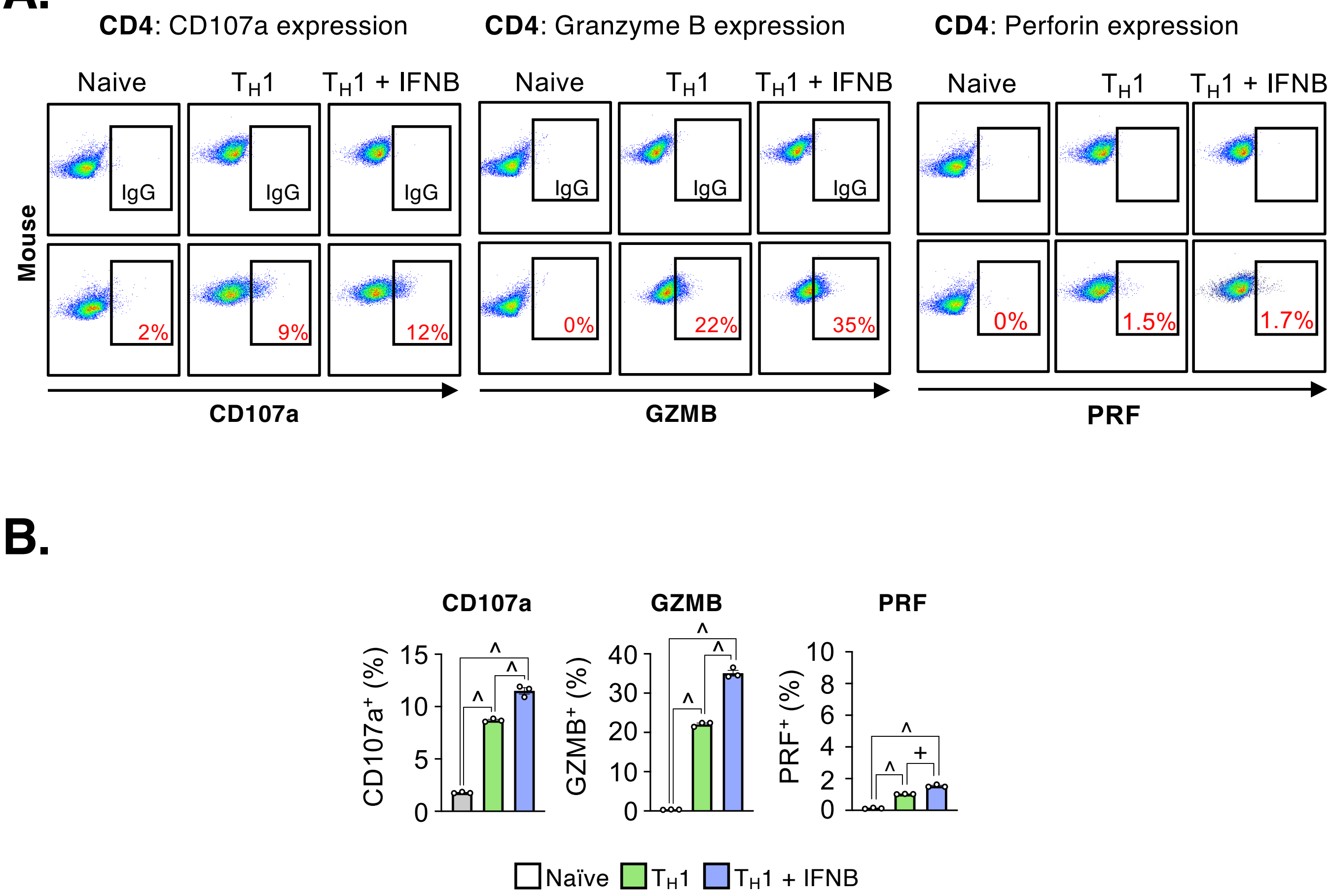

CD4: CD107a expression
CD4: Granzyme B expression
CD4: Perforin expression
Naive
TH1
TH1 + IFNB
Mouse
IgG
2%
9%
12%
0%
22%
35%
0%
1.5%
1.7%
CD107a
GZMB
PRF
CD107a
GZMB
PRF
CD107a+ (%)
GZMB+ (%)
PRF+ (%)
^
+
Naïve
TH1
TH1 + IFNB

**Figure S7. IFNB induces increased degranulation and expression of GZMB and PRF in murine CD4$^+$ T-cells.** (**A**). Representative flow cytometry plots showing cell surface CD107a and intra-cellular GZMB and PRF in naïve or $T_H$1-activated CD4$^+$ T-cells alone or following treatment with IFNB. (**B**) Quantification of the proportion of CD107a, GZMB and PRF positive CD4$^+$ T-cells following the treatments outlines in 'A' (n = 3 per condition).

### Figure S8

**A.**

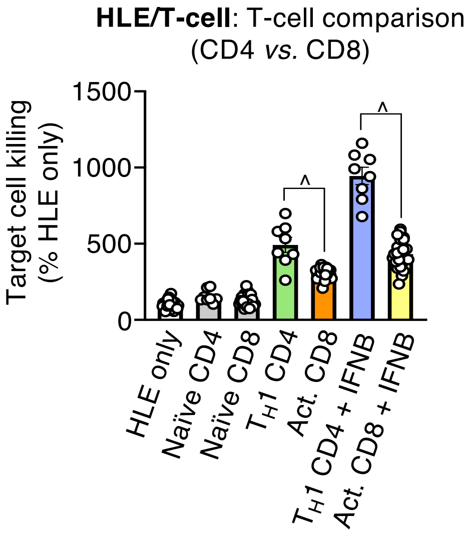


**B.**

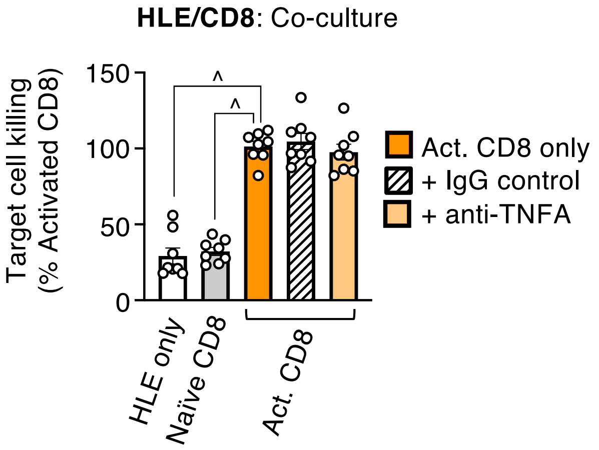


**C.**

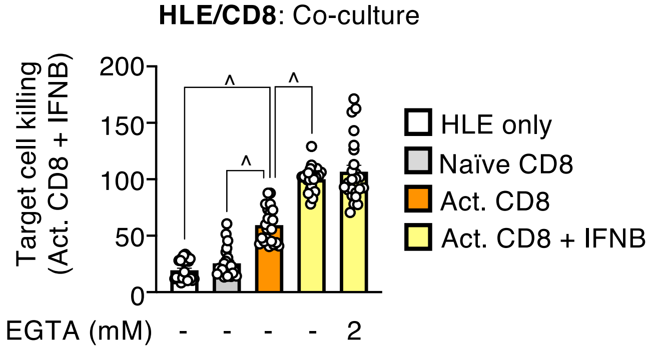

**Figure S8. CD8$^+$ T-cells display inferior tumoricidal activity against HCC cells compared with CD4$^+$ $T_H$-cells and use neither TNFA nor degranulation as their principal mode of killing. (A)** Co-culture assay between human HLE HCC cells and either naive or activated T-cells in the presence of absence of IFNB (500 ng/mL). (**B**) Co-culture assay between human HLE HCC cells and naive or activated CD8$^+$ T-cells in the presence of anti-human TNFA neutralizing antibody (10 μg/mL) or isotype matched control. (**C**) Co-culture assay between human HLE HCC cells and either naive or activated CD8$^+$ T-cells with or without IFNB (500 ng/mL) in the presence of the degranulation inhibitor 'EGTA' (2 mM). The degree of target cell killing was determined in all co-culture assay by flow cytometry following incubation overnight. (Act. = activated, Sora. = sorafenib; * $p<0.05$, ≠ $p<0.01$, + <0.001, ^ <0.0001; n = 8 per condition).
